# In Vitro Modulation of MECP2 Expression via Antisense Inhibition of miR-132-3p in SH-SY5Y Cells

**DOI:** 10.1101/2025.10.06.680678

**Authors:** Arda Orcen, Beyza Goncu, Baris Ekici, Burak Tatli, Emrah Yucesan

## Abstract

Rett syndrome (RTT) is a severe neurodevelopmental disorder that predominantly affects females. It is characterized by developmental regression during infancy, including loss of speech, gait abnormalities, intellectual disability, seizures, respiratory issues, and stereotypic hand movements. RTT is mainly caused by spontaneous, non-inherited mutations in the methyl-CpG-binding protein 2 (MECP2) gene located on the X chromosome. Despite considerable scientific progress, no effective treatments currently exist for MECP2-related pathology. Antisense oligonucleotides (ASOs), which selectively bind and inhibit specific RNA targets, have recently emerged as promising therapeutic agents. This study investigated the effects of ASO-mediated inhibition of miR132-3p on MECP2 mRNA expression in the SH-SY5Y neuroblastoma cell line to evaluate its therapeutic potential in RTT. Our results demonstrated a significant reduction in miR132-3p expression 6 hours after transfection with Mixmer and 2’OMe-modified ASOs. Correspondingly, MECP2 mRNA levels were significantly upregulated in all ASO-treated groups, with the most prominent increase observed at 12 hours post-transfection in the 2’OMe group. This time-dependent inverse relationship between miR132-3p and MECP2 expression supports the regulatory interaction between them. These findings suggest that 2’OMe-modified ASOs targeting miR132-3p may represent a promising therapeutic strategy for RTT and warrant further in vivo investigation.

## Introduction

Rett syndrome (RTT) is a severe X-linked neurodevelopmental disorder that primarily affects females and is characterized by developmental regression in early childhood (Kaufmann et al. 2016). Clinical features include loss of purposeful hand skills and spoken language, gait abnormalities, intellectual disability, seizures, breathing irregularities, and stereotypic hand movements (Percy et al. 2023). RTT is clinically classified into classic and variant forms. Classic RTT includes all four main diagnostic criteria, while variant RTT presents with atypical features and is further subcategorized into preserved speech, early-onset, and congenital variants (Neul et al. 2010). RTT predominantly affects females and is caused by de novo mutations in the MECP2 gene located on the X chromosome, which are usually not inherited (Petriti et al. 2023). The treatment process for RTT involves a multidisciplinary approach (Vilvarajan et al. 2023). Although no curative treatment is currently available, various therapeutic approaches are under investigation, ranging from physiotherapy (e.g., applied behavior analysis, conductive education, environmental enrichment, hydrotherapy, and music therapy) to advanced gene-editing technologies (Fonzo et al. 2020; Panayotis et al. 2023). While studies on gene therapy are limited, clinical applications remain at an early stage (Sandweiss et al. 2020). Another promising approach involves antisense oligonucleotides (ASOs), designed to regulate MECP2 levels—a key factor in RTT pathology (Sztainberg et al. 2015; Shao et al. 2021).

ASOs are chemically modified nucleic acid sequences capable of binding target mRNAs or microRNAs with high specificity. Several ASO-based therapies are currently under investigation for neurological diseases (Brunet de Courssou et al. 2022; Sardone et al. 2017; Bennett et al. 2019; Scoles and Pulst 2019). In light of this information, this study aimed to investigate whether ASOs targeting miR132-3p, a microRNA involved in MECP2 regulation, could upregulate MECP2 mRNA expression in vitro, thereby highlighting a possible therapeutic avenue for RTT.

## Materials and Methods

### ASO Design

Three chemically modified ASOs targeting miR132-3p were synthesized to optimize stability, binding affinity, and therapeutic potential:

#### 1. Locked Nucleic Acid (LNA)

Enhances binding affinity by structurally constraining the ribose sugar backbone, increasing hybridization efficiency and nuclease resistance.

#### 2. 2’-O-Methyl (2’OMe)

Offers high molecular stability and reduced immunogenicity with moderate affinity.

#### 3. Mixmer

Combines LNA and 2’OMe modifications to balance stability, binding affinity, and safety.

All ASOs were designed to inhibit miR132-3p and influence MECP2 expression in SH-SY5Y cells.

### Cell Culture and Synchronization

SH-SY5Y neuroblastoma cells (ATCC, CRL-2266) were cultured in DMEM/F12 medium (1:1) supplemented with 10% Fetal Bovine Serum (FBS), 1% penicillin-streptomycin, and 0.1% primocin maintained at 37°C in a 5% CO_2_ incubator. Cells were passaged every 2–3 days, and experiments were performed using cells between passage 5–9. To synchronize the cell cycle, cells were serum-starved in 3% FBS for 24 hours to induce G0 phase arrest, followed by re-stimulation with 10% FBS for 24 hours before transfection (Dravid et al. 2021).

### ASO Transfection

Cells were transfected with 10 nM of each ASO using Lipofectamine RNAiMax in OptiMEM medium. Transfection durations were 6, 12, and 24 hours. Additional groups were cultured for 24 hours post-transfection to assess ASO sustainability. Transfection efficiency was indirectly assessed through analysis of cell viability and morphological integrity (Muse Cell Analyzer (Merck Millipore)).

### Total RNA Isolation and cDNA Synthesis

Total RNA was isolated using the EcoPURE Total RNA Kit (Ecotech, Erzurum, Türkiye). RNA quantity and quality were determined by 260/280 nm absorbance ratios. cDNA synthesis was carried out with specific kits for mRNA and miRNA conversion (Applied Biological Materials Inc., Canada).

### qRT-PCR

*MECP2* gene primers were designed by using Primer3Plus and validated via In-Silico PCR (UCSC), and BLAST (NIH). The optimal melting temperature was determined by gradient PCR in a T100 thermal cycler (Bio-Rad, CA, USA) and analysed by agarose gel electrophoresis. *GAPDH* was used as a reference control for mRNA expression levels. For miR132-3p gene, primer was purchased from ABM (cat no: MPH02174) and as reference control miR-let-7a (ABM, cat no: MPH02041) were used. All cDNA samples were diluted for 1:5 with ultrapure water before further experiments. After that, the expression levels were evaluated with BlasTaq™ 2X qPCR MasterMix (Applied Biological Materials Inc. (ABM), BC, Canada) according to the manufacturer’s instructions with the CFX Connect Real-Time PCR Detection System (Bio-Rad, CA, USA). Samples were tested as duplicates on the same 96-well PCR plate per sample in 40 cycles, and the mean Cq values were used for further analysis. RT-qPCR was performed with a two-step method including 95°C for 2 minutes, then denaturing at 95°C for 10 seconds, annealing at 60°C for 10 seconds, and extension at 72°C for 10 seconds.

## Results

The experimental procedures commenced with the culture of SH-SY5Y cells until a confluency greater than 70% was achieved. To synchronize the cell cycle, a serum starvation step was implemented by reducing the fetal bovine serum (FBS) concentration to 3% for 24 hours. Subsequently, cells were reintroduced into 10% FBS medium for 24 hours prior to transfection (Figure 1A). This synchronization strategy resulted in a marked increase in the proportion of cells in the G0/G1 phase, rising from 47.6% to 68.9%, thereby producing a more homogeneous population. Synchronization aimed to enhance transfection efficiency by promoting a uniform cell cycle stage and, consequently, consistent ASO uptake.

**Figure 1.**
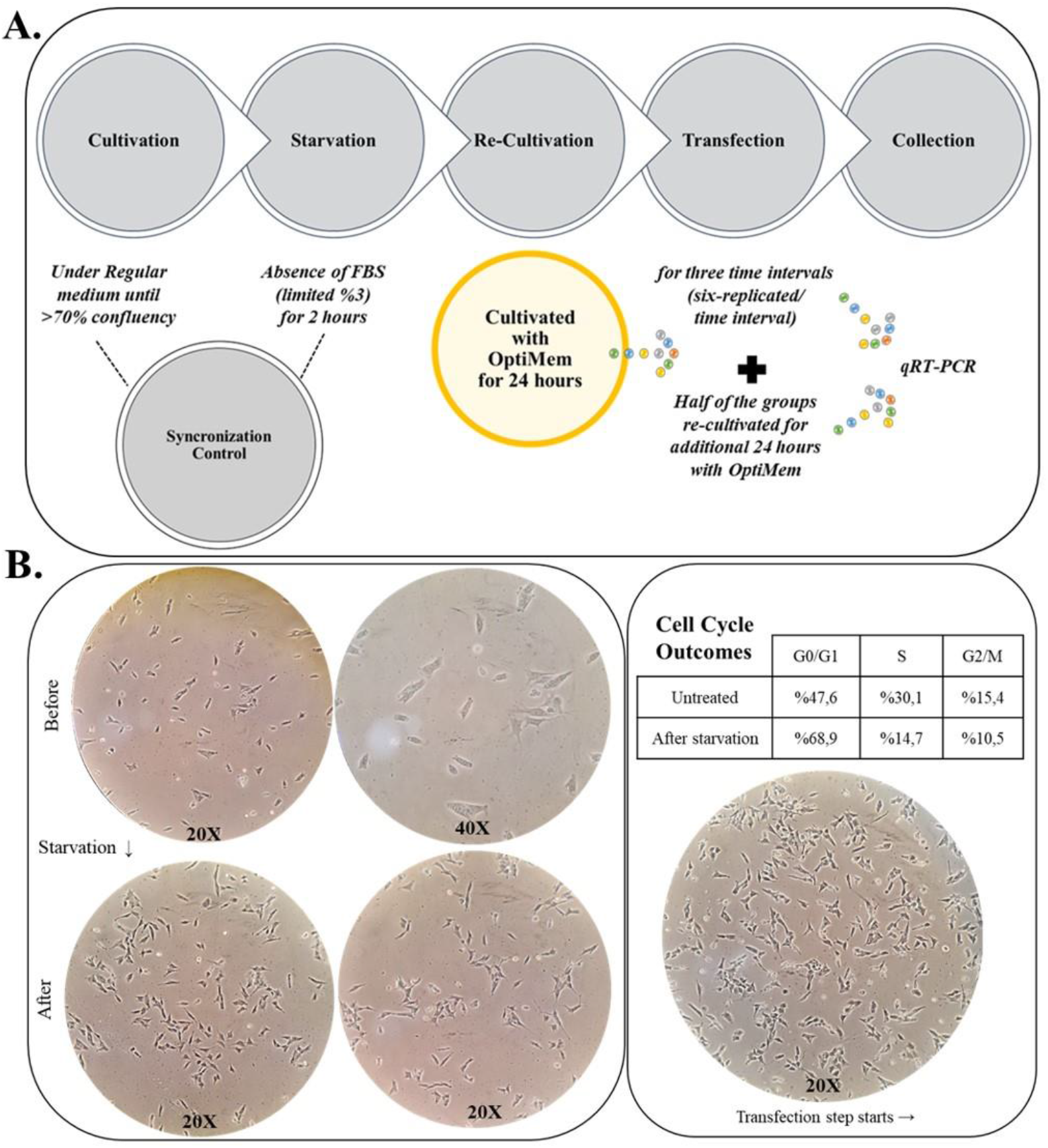
**A)** The synopsis of the experimental procedures that were employed in the study with SH-SY5Y cells. **B)** Demonstrates the transfection and morphological follow-up stages prior to and following cell cycle synchronization of SH-SY5Y neuroblastoma cells. Cell cycle analysis was performed to determine the number of cells in the G0/G1 phase, with the aim of achieving a homogeneous cell population for transfection.

Morphological observations (Figure 1B) confirmed that cells maintained structural integrity and viability throughout the experimental timeline. Cell viability assays were performed in parallel, with six biological replicates monitored at each of the 6-, 12-, and 24-hour time points following transfection. To evaluate the persistence of ASO effects, half of the experimental groups were further cultured for an additional 24 hours post-transfection, after which total RNA was extracted for downstream molecular analyses.

Quantitative real-time PCR (qRT-PCR) analysis revealed a statistically significant downregulation of miR132-3p expression at 6 hours post-transfection with both Mixmer and 2’OMe-modified ASOs (p = 0.0368 and p = 0.0086, respectively), compared to the negative control group (Figure 2A). At 12 hours post-transfection, a non-significant but observable downward trend in miR132-3p expression persisted across all ASO-treated groups, including the LNA group. By 24 hours, expression levels exhibited more variability, with no consistent trend across the different ASO formulations, suggesting potential degradation or reduced activity over time.

**Figure 2.**
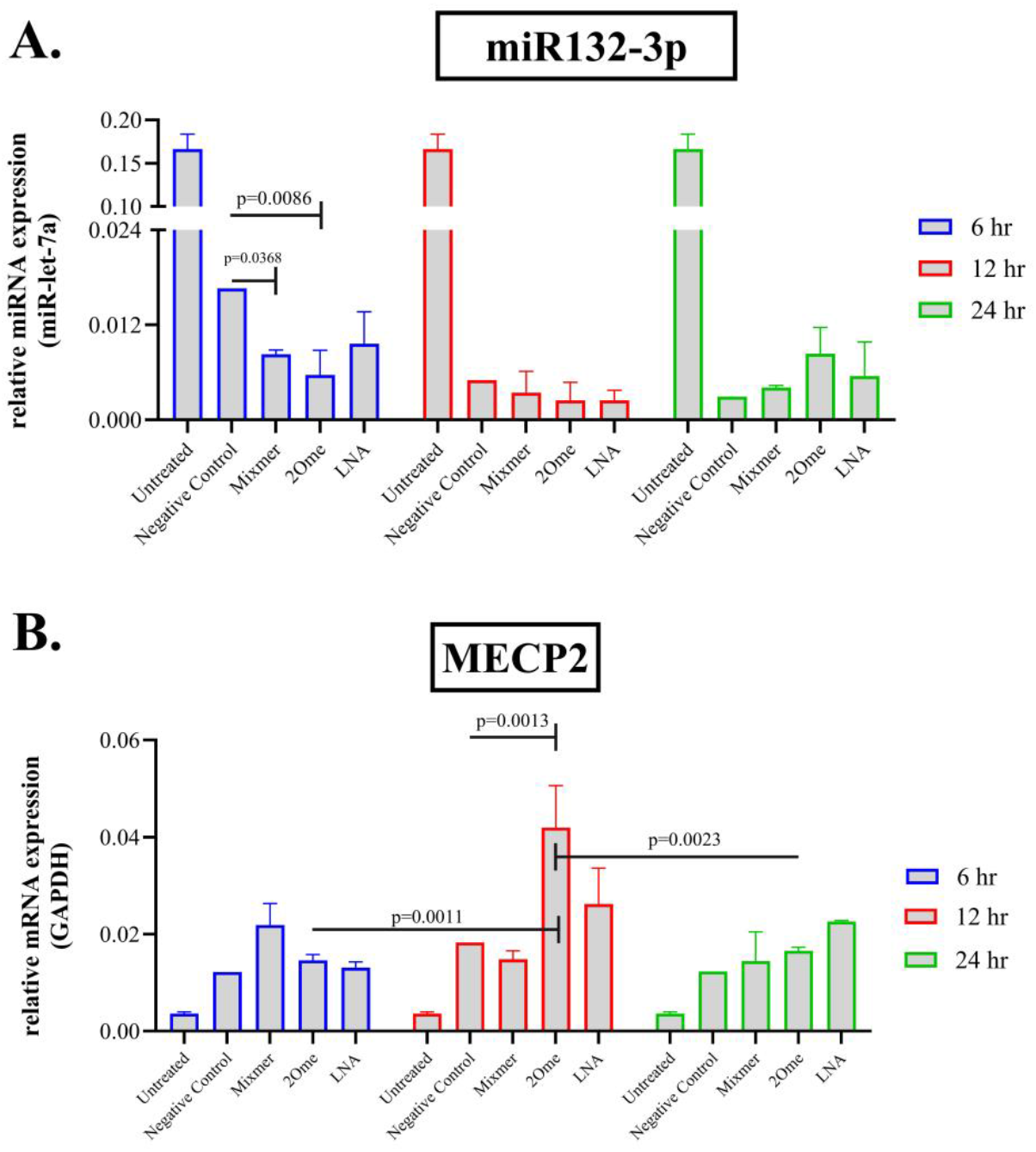
Changes in miR132-3p and MECP2 Expression with ASO Transfection. **A)** Transfection outcome at three different time intervals. miR-let-7a was used as the reference miRNA. **B)** MECP2 mRNA expression levels after transfection. A significant increase observed (p < 0.0001) in all ASO treatment groups, GAPDH was utilized as the reference gene.

In parallel, MECP2 mRNA expression levels were quantified to determine the functional consequence of miR132-3p inhibition. Notably, all ASO-treated groups exhibited a statistically significant increase in MECP2 mRNA compared to the untreated control (p < 0.0001). Among the ASOs tested, the 2’OMe-modified oligonucleotide demonstrated the most robust and consistent upregulation. At 12 hours post-transfection, MECP2 expression reached its peak in the 2’OMe group, showing a significant increase compared to both the 6-hour and 24-hour time points (Figure 2B). This peak expression coincided with the earlier downregulation of miR132-3p, particularly evident after 6 hours, suggesting a temporal regulatory relationship between the two molecules.

These findings indicate that 2’OMe-modified ASOs exert the most pronounced and sustained effects on miR132-3p suppression and subsequent MECP2 mRNA upregulation. These time-dependent effects highlight the importance of kinetic analysis for optimizing ASO-based therapeutic strategies.

## Discussion

ASO treatment options are considered a highly promising approach today—not only as a driving force behind numerous current studies, particularly due to advancements in technology and the diversification of chemical applications, but also because of their potential to lead to concrete treatment alternatives in the future. Numerous studies in the literature on ASO therapies have found opportunities for application across a wide spectrum, ranging from cancer to neurology. The clinical application of ASO therapies began with the FDA approval of Fomivirsen in 1998 (Dhuri et al. 2020). Following this pioneering drug, Mipomersen was introduced in 2016 for the treatment of Familial Hypercholesterolemia (Thomas et al. 2013), and shortly thereafter, Eteplirsen was presented for Duchenne muscular dystrophy (DMD) (Mendell et al. 2016). This progress has spurred numerous subsequent studies targeting a variety of neurological diseases—from rare pediatric disorders to common neurodegenerative conditions—using similar ASO-based approaches (Bennett et al. 2019; Li et al. 2021; McDowall et al. 2024; Zhang 2024).

Within this context, miR-132-3p, the focus of our study, has been reported to play a role in stress response through evolutionarily conserved pathways (Haviv et al. 2018). Initially identified in mouse neural tissues, miR-132 has since been detected in humans, zebrafish, and cows, illustrating its evolutionary conservation (Lagos-Quintana et al. 2001; Calin et al. 2004; Chen et al. 2005; Coutinho et al. 2007). Functionally, miR-132-3p has been implicated in the etiopathogenesis of neurological disorders such as Alzheimer’s disease and chronic neuropathic pain (Leinders et al. 2016; Zeng et al. 2022), and it exerts broad effects across the nervous system by modulating neuronal processes like axon growth, migration, and neuroplasticity (Qian et al. 2017; Mu et al. 2024). Given its multifaceted role in neuronal development and disease, miR-132-3p represents a compelling target for ASO-based interventions.

On the other hand, Methyl CpG binding protein 2 (MeCP2), a key factor in the pathogenesis of Rett syndrome (RTT), regulates chromatin structure via binding methylated DNA (Liyanage and Rastegar 2014; Martínez de Paz and Ausió 2017). The MECP2 gene is essential for normal brain development, and its dysfunction leads to neurodevelopmental abnormalities (Kaufmann et al. 2005). Studies in Mecp2-deficient mouse models have revealed dysregulation of numerous miRNAs (65 out of 245 analyzed) show altered expression patterns, indicating complex miRNA-mediated regulatory networks influenced by MeCP2 (Urdinguio et al. 2010). Further research using human iPSC models demonstrated that MeCP2 modulates neurogenesis through novel miRNA pathways, including miR-197-mediated inhibition of the NOTCH pathway during neuronal progenitor cell differentiation (Mellios et al. 2018; Wang et al. 2019). Taken together, these findings highlight the critical role of MECP2 and its associated miRNAs in neurological disease pathogenesis.

Specifically relating to miR-132 and RTT, studies examining the MECP2 isoforms, BDNF, and miR-132 levels across different brain regions suggest region-specific regulation of the MECP2E1/E2-BDNF-miR132 axis (Pejhan et al. 2020). Moreover, in clinical samples from RTT patients, only miR-483-5p and miR-132-3p among several significant miRNAs showed increased expression, emphasizing miR-132-3p’s involvement in RTT pathology (Castells et al. 2021).

Considering these lines of evidence, miR-132-3p represents a promising therapeutic target for RTT. Recent investigations into antisense oligonucleotides (ASOs) designed to inhibit miR-132-3p provide compelling data: chemically modified ASOs, such as those incorporating 2’-O-Methyl (2’OMe) and Mixmer modifications, effectively downregulate miR-132-3p and concomitantly upregulate MECP2 mRNA, underscoring their potential for clinical application in restoring MECP2 function.

Our study corroborated these findings, demonstrating that 2’OMe ASOs elicited rapid and robust downregulation of miR-132-3p within 6 hours of transfection, followed by significant MECP2 upregulation at 12 hours. This temporal sequence suggests a direct mechanistic link between miRNA inhibition and target mRNA stabilization. Additionally, cell cycle synchronization of SH-SY5Y cells in G0/G1 phase prior to transfection significantly enhanced ASO uptake and efficacy, consistent with previous reports on improved transfection efficiency under synchronized conditions (Wu et al. 2005; Kevadiya et al. 2023).

However, the observed variability in miR-132-3p suppression and MECP2 expression at 24 hours points to potential degradation or diminished ASO activity over time, warranting further pharmacokinetic and stability studies in vitro and in vivo. Given the multifactorial nature of RTT pathogenesis, future research should also explore combination therapies targeting multiple dysregulated miRNAs or signaling pathways to enhance therapeutic outcomes.

Finally, while ASO-based approaches offer promising avenues for RTT treatment, their clinical translation requires addressing challenges related to safety and specificity. Immune activation and off-target effects remain major concerns, as highlighted by Pradère et al. (Gagliardi and Ashizawa 2021). The development of advanced chemical modifications and targeted delivery methods will be critical to overcoming these hurdles and enabling effective, safe clinical applications (Hammond et al. 2021; Zhu et al. 2022).

In conclusion, this study provides strong evidence supporting ASOs targeting miR-132-3p as viable therapeutic agents in RTT. The pronounced effects of 2’OMe ASOs on miR-132-3p downregulation and MECP2 upregulation underscore their therapeutic potential and set the stage for future preclinical and clinical development toward effective RTT treatments.

## Acknowledgments

We sincerely thank the individuals diagnosed with Rare Disease in general and Rett Syndrome in particular, who inspired this study.

